# What happens on islands, doesn’t stay on islands: Patterns of synchronicity in mosquito nuisance and host-seeking activity between a mangrove island and adjacent coastal development

**DOI:** 10.1101/2020.04.21.054262

**Authors:** Brian J. Johnson, Russell Manby, Gregor J. Devine

**Author notes:** Corresponding author at: QIMR Berghofer Medical Research Institute, 300 Herston Road, Herston, QLD 4006, Australia, (B. J. Johnson).

## Abstract

Coastal development is expanding globally in response to mass human migration, yet urban planning guidelines often overlook the problems that human encroachment on or near coastal mosquito habitat may cause. This study aimed at elucidating the frequency and magnitude of dispersal of highly vagile saltmarsh mosquitoes from productive off-shore bay islands to adjacent coastal human developments. Inter-population dynamics and daily host-seeking activity of saltmarsh mosquitoes were monitored daily at 15-minute intervals within a productive bay island and adjacent coastal development in southeast Queensland, Australia, using emerging smart trap technology over a 2-month period of high mosquito activity. The regulation of mosquito dispersal and host-seeking activity by local environmental factors, e.g. temperature, relative humidity and hourly wind patterns, were also investigated. The data show that the primary saltmarsh mosquitoes *Aedes vigilax* (Skuse) and *Culex sitiens* (Wiedemann) disperse from offshore breeding sites to neighboring mainland areas in high numbers and in highly synchronized waves despite unfavorable wind patterns and the need to traverse a considerable expanse (ca. 1.4 km) of open water. Patterns of host-seeking activity within each site were also remarkably similar despite notable differences in the local environment demonstrating a consistency in host-seeking activity across disparate habitats. These findings demonstrate that distant saltmarsh habitats, including offshore breeding sites, are likely to be primary sources of mosquito nuisance for coastal housing developments. This observation highlights the need to develop new planning and regulatory guidelines that alert urban planners to the risks of encroaching on habitats close to the sources of highly vagile mosquito species.

## 1. Introduction

The development and transformation of coastal zones has greatly increased during recent decades to accommodate mass human migration to these areas— a trend which is expected to continue in the future (Neumann et al., 2015; Seto, 2011; Seto et al., 2011). Australia is not exempt from this trend and > 85% of Australians now live within 50 km of the coast (ABS, 2016). This is particularly evident around southeast Australia’s metropolitan centers where the agreeable climate and access to coastal recreational activities are highly attractive (Stimson and Minnery, 1998). The emphasis on continued growth in this region has resulted in the encroachment of human dwellings into marginal coastal wetlands and saltmarshes areas (Dwyer et al., 2016; Ramasamy and Surendran, 2012). This encroachment has resulted in a predictable increase in complaints of biting nuisance by the residents of these new developments (Ryan et al., 2006). Those rate payers expect the regions’ local government mosquito control programs to mitigate the problem. This political pressure increases the burden on existing control programs. Local and state governments attempt to address mosquito issues by implementing control measures, planning strategies and development guidelines that aim to reduce contact with mosquitoes and while adhering to environmental law and the need to conserve coastal ecosystems (Dwyer et al., 2016; Russell, 1998; Tomerini et al., 2011). The majority of efforts and resources are allocated to area-wide larval control programs and the physical management of natural habitats to reduce mosquito productivity (Dale and Breitfuss, 2009; Dale and Knight, 2006; Russell and Kay, 2004). Habitat management has principally been applied to mainland features, and, while there have been successes (Dale and Breitfuss, 2009; Dale and Knight, 2006), highly productive offshore sources of mosquitoes remain problematic and pose particularly intractable control problems for neighbouring developments on the mainland.

In the Australian southeast, offshore sources of biting insects are predominantly uninhabited bay islands dominated by a mixture of mangrove and saltmarsh habitat highly favorable to *Aedes vigilax* (Skuse) mosquitoes (Dale and Knight, 2006; Griffin et al., 2010; Knight et al., 2012). This and other saltmarsh-associated species represent significant health threats as they are important vectors of Ross River virus (RRV) and Barmah Forest virus (BFV), the two most common mosquito-borne viruses in Australia (Jacups et al., 2008; Naish et al., 2011; Ryan et al., 2006; Tomerini et al., 2011). Of the two diseases, RRV is the most medically important with close to 5000 cases reported each year in Australia. The human disease manifestations of RRV infection are consistent with other alphaviruses and include febrile illness, polyarthritis, and rash (Harley et al., 2001). BFV has similar symptoms to RRV but is less prevalent (Naish et al., 2011). Due to the combined health and nuisance risks that *Ae vigilax* presents, the majority of state and local funding for mosquito control is focused on controlling its abundance through area-wide, aerial larviciding campaigns (Russell et al., 2003). In the state of Queensland, operational costs currently exceed AUD 20 million annually, increasing from AUD 7 million in 1993 and AUD 11 million in 2004 (Tomerini, 2007; Tomerini et al., 2011). Although such programs can be highly effective (Becker, 2006; Floore, 2006; Lacey, 2007), they are operating in conditions that make it difficult to attain perfect control (e.g. heavy vegetation cover prevents penetration of larvicides and flushing tides dilute product). Some production of adult mosquitoes is therefore inevitable, but few programs have contingencies in place to control adult nuisance in residential areas. Adulticiding is rarely used option as it is a costly and fleeting form of control with a great deal of public resistance over non-target impacts and widely-publicized misuses by reckless operators in other countries (Bonds, 2012; Boyce et al., 2007; Oberhauser et al., 2009). Programs that do have the capacity to control adults do so mostly in cases of extreme nuisance in response to residential complaints or heightened disease activity (Tomerini, 2007). Consequently, vagile mosquito populations originating from uninhabited but highly productive bay islands represent a significant nuisance and public health threat to adjacent mainland developments and a political headache for regional mosquito control programs.

Amongst mosquitoes, the dispersal capabilities of saltmarsh *Aedes* mosquitoes are unrivalled (Provost, 1960; Service, 1997). The dispersal distances of these species are often so great that direct quantification from traditional mark-release-recapture experiments is impossible due to exponentially diminishing probabilities of insect recapture over large distances (Jardine et al., 2014). In the case of *Ae. vigilax*, a recent quantitative report confirmed dispersal distances of at least 3 km from coastal breeding sites, the distance limit of the sampling scheme (Webb and Russell, 2019). This observation is supplemented by numerous qualitative reports and population genetic studies that report dispersal over much greater distances (Chapman et al., 1999; Webb and Russell, 2019). Although few reports exist for other prominent saltmarsh species such as *Culex sitiens* (Wiedemann), reports on sibling species indicate that long-distance dispersal is common amongst saltmarsh species (Bryan et al., 1992; Russell, 1986; Webb and Russell, 2019). This vagility probably places a large proportion of coastal residents within the reach of mosquito populations originating from distant breeding sites but there are few empirical studies on the timing and magnitude of these dispersal events.

The difficulty in determining the nature and magnitude of mosquito dispersal from offshore breeding sites to mainland human developments arises from a problem of logistics. These areas are often remote, and any evaluation of mosquito productivity or treatment efficacy requires boat access. The routine use of traditional population surveillance tools, such as battery-powered, CO_2_-baited light traps are therefore impractical for intensive, longitudinal surveillance of daily activity and abundance (Ritchie and Kline, 1995). Although there are traps that allow for multiple collections over a period of days or even weeks, they have a limited number of collection chambers resulting in poor temporal resolution and require frequent servicing (Barnard et al., 2011). Mark-release-recapture experiments have long been the gold-standard for elucidating dispersal capability (Guerra et al., 2014), but the recapture of highly dispersive mosquitoes is extremely challenging. Large numbers of traps must be serviced almost daily over large distances and huge numbers of mosquitoes must be sourced, marked and released (Guerra et al., 2014; Webb and Russell, 2019). Thus, these studies are often performed over short periods with little to no replication. This, combined with extremely low recapture rates (in same cases <1% (Webb and Russell, 2019), greatly limits the usefulness of this method for long dispersing mosquitoes like *Ae. vigilax* and *Cx. sitiens*.

Usefully, great advances have recently been made in the development of ‘smart’ mosquito traps. Smart traps employ the use of one or more sensors to optimize trapping efficiency. In some cases the traps can remotely collect and transmit this information for the purposes of vector surveillance (Fernandes et al., 2018). Many of the current designs incorporate optical sensors, mostly infrared-based light arrays, which detect small variations in the light captured by phototransistors as the insect enters the trap (Batista et al., 2011; Potamitis and Rigakis, 2015). These variations can then be used to differentiate species and sexes based on their unique wing-beat frequencies or body size. The BG-Counter (Biogents AG, Regensburg, Germany) is the first commercially available smart trap to incorporate this technology. It has some capacity to automatically differentiate mosquitoes from other insects entering the trap, count them, and wirelessly transmit the results (Fanioudakis et al., 2018). This technology has begun to be adopted by control and surveillance agencies (Lucas et al., 2019) and makes quantifying fine-scale population dynamics and aspects of host-seeking activity feasible within the most logistically challenging environments.

The objective of this study was to adopt emerging smart trap technology to investigate patterns of synchronicity in saltmarsh mosquito abundance and host-seeking activity between a highly productive offshore breeding site and an adjacent coastal development devoid of notable breeding area in southeast Queensland, Australia. This empirical information can then be used to illustrate the hazards of building new developments in close proximity to extremely productive mosquito habitats and the importance of designing sustainable strategies to mitigate nuisance and disease risk to coastal residents.

## 2. Methods

### 2.1 Study Sites

The study occurred in the area of Redland Bay, Queensland, Australia from April to June 2019. Redland Bay is a coastal locality located 35 km south-east of Brisbane, the capital of Queensland. The bay, part of Southern Moreton Bay, contains a number of inhabited and uninhabited islands (Fig. 1 A). The uninhabited islands contain a combined 359.7 ha of productive saltmarsh breeding habitat for both *Ae. vigilax* and *Cx. sitiens.* The average distance from the islands to the coast is 2.82 km (0.48-5.4 km), a distance well within the reported and estimated dispersal ranges of both species (Fig. 1 B) (Bryan et al., 1992; Webb and Russell, 2019). To investigate the possibility of dispersal of bay island populations to mainland Redland Bay, daily mosquito abundance and host-seeking activity was monitored within a coastal residence and an adjacent uninhabited bay island (Fig. 1 C). The selected bay island contains 112.2 ha of saltmarsh mosquito breeding habitat and is the closest of the islands (ca. 1.4 km edge to edge) to the selected residential area. The island is known to produce vast numbers of *Ae. vigilax* and *Cx. sitiens* mosquitoes (Kay and Jorgensen 1986 and Chapman et al. 1999) and is dominated by *Sporobolus virginicusis* (Saltwater couch), *Sarcocornia quinqueflora* (Bead weed) and *Avicennia marina* (Grey mangrove) vegetation. The mainland residential site is located approximately northwest of the island in the suburb of Sandy Cove. The suburb of Sandy Cove has produced numerous resident nuisance complaints for saltmarsh mosquitoes over the years (Ryan et al. 2006) despite being in an area devoid of notable saltmarsh mosquito breeding habitat due to extensive residential and agricultural development.

**Figure 1.**
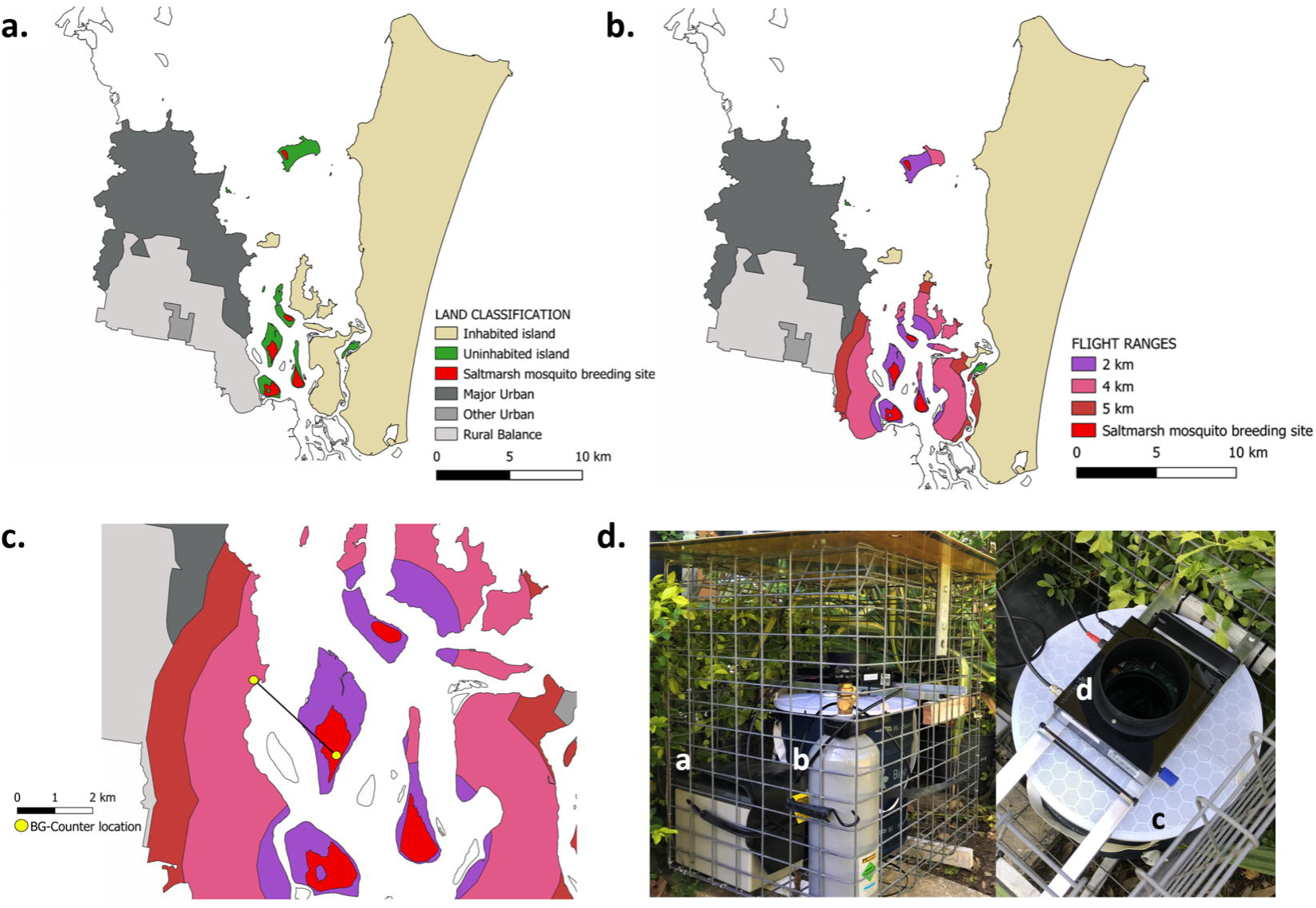
(A) Land classifications of Redland City located in southeast Queensland, Australia. (B) Area at risk of mosquito nuisance originating from uninhabited bay islands across a range of observed saltmarsh mosquito dispersal distances (2, 4 and 5 km). (C) Adult mosquito monitoring locations. Depicted flight ranges are based on previous reports from mark-release-recapture experiments (Bryan et al., 1992; Chapman et al., 1999; Webb and Russell, 2019). (D) The BG-Counter trap system used in the study highlighting the setting of the BG-Counter unit on top of the BG-Sentinel trap. The trap components shown include: (a) 12 V rechargeable battery (70 Ah), (b) CO_2_ gas cylinder (2 kg), (c) BG-Sentinel trap body, and (d) BG-Counter unit.

### 2.2 Trap Operation and Climate Monitoring

Daily population and host-seeking activity data were obtained through the use of BG-Counter (Biogents AG, Regensburg, Germany) units in combination with the BG-Sentinel trap (Maciel-de-Freitas et al., 2006) to create a complete trapping system (Fig. 1D). The units were set to record collection totals every 15 min allowing us to generate fine-scale reports of biting activity. The traps were baited with CO_2_ at a rate of 200 ml/ min provided from 2.5 or 6.0 kg gas cylinders using the supplied regulator. The different sized gas cylinders allowed for run times of 5 days and 12 days, respectively. Power was supplied by a 70 Ah 12 V rechargeable lead acid battery (HGL70-12, Fullriver Battery, Camarillo, CA USA) on the bay island site and by either battery power or household (mains) power in the residential site. The 70 Ah batteries produced 140 run time hours powering both the counter unit and trap fan. To prolong battery life on the bay island, a 40 W solar panel was connected that allowed for battery reset intervals of 3-4 weeks. The BG-Sentinel trap in the residential site was set as specified by the manufacturers with the provided collection bags as the capacity of the bags was adequate for the numbers captured. Due to the large collection totals on the island, the standard collection bags could not be used. The trap was instead run without any inner collection bags, the insects being collected in the body of the BG-Sentinel trap. The inside of the main trap compartment was treated with insecticide surface spray (Mortein Outdoor Barrier Surface Spray; imiprothrin 0.3 g/kg and deltamethrin 0.6 g/kg) to minimize the number of escapees and reduce counting errors due to escapees. The timing of fan and counter operation, as well as CO_2_ output, was controlled through the BG-Counter web application. The BG-Counter contains environmental sensors that enable the recording of temperature, relative humidity and ambient light. All three variables were recorded daily at 15 min intervals. Daily wind speed and direction data (Fig. 2) were obtained through the Bureau of Meteorology (www.bom.gov.au) for the Redland Station (−27.5433, 153.2394). All trap components and CO_2_ cylinders were placed in purpose-built cages (galvanized rolled-top steel; 75 mm^2^ openings) to protect components from theft and damage.

**Figure 2.**
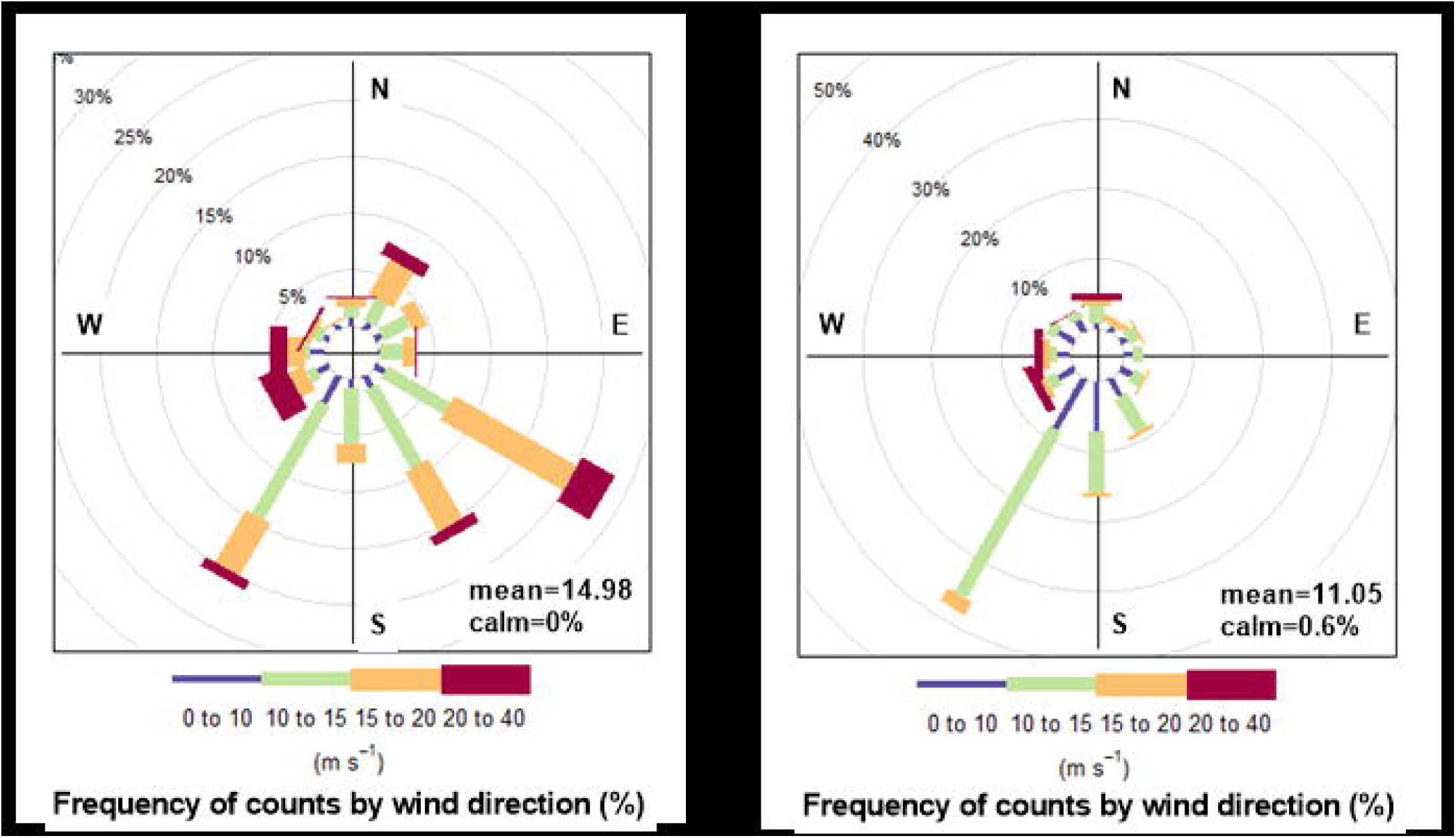
(A) Daytime (6 am-6pm) and nighttime (6 pm-6 am) hourly wind speed and direction summaries for Redland City during the study period. Wind direction is reported as the direction from which the wind originated (i.e., wind would be blowing in opposite direction).

### 2.3 Mosquito Collection and Identification

Captured mosquitoes were collected at various intervals ranging from 2-6 d for the residential site and 6-15 d for the bay island. The collections were brought back to the council depot and stored at −20 C until identified. Subsamples of each collection were identified morphologically using standard keys (Ehlers and Alsemgeest 2010; Marks 1982) to generate biting insect community compositions and to determine the level of agreement between manual counts and those produced from the BG-Counter.

### 2.4 Data Analysis

Paired *t-*tests were used to evaluate differences in daily mosquito abundance between the residential and saltmarsh sites. Analyses were done for both total daily collections and for estimated daily collections of *Ae. vigilax* and *Cx. sitiens.* Estimated abundances were determined by multiplying daily collection totals by the community dominance (%) of each species observed during each surveillance interval. One-way ANOVA was performed to test for differences in biting activity at 3-hr intervals within a site and two-way ANOVA was performed to test for differences in biting activity between sites. We used generalized least squares linear regression to compare trends in mosquito populations between the two sites with time (day) included as a first-order autoregressive (AR1) term. The use of the AR1 term was confirmed by analysis of autocorrelation plots. Additional explanatory variables included wind speed, wind direction, and mean daily temperature. All analyses involving mosquito counts were performed on log + 1 transformed data and percentage data were subjected to arcsine transformation prior to analysis.

## 3. Results

### 3.1 Mosquito Collections

The traps were operated for a total of 50 trap days from April 2^nd^ to May 20^th^, 2019. During this period, mosquitoes captured at the residential and bay island sites were collected and identified 13 and 6 times, respectively. *Culex sitiens* and *Ae. vigilax* were the dominant species within both sites, accounting for 95.09 ± 2.78% of residential and 99.0 ± 0.97% (Fig. 3A) of bay island collections (Fig. 3B). Daily collection totals of these species were significantly greater (*t*_49_=20.32, *p*<0.001) in the bay island site (1551 ± 227.7 mosquitoes/day) compared to the residential site (194.9 ± 20.87), a trend that was observed throughout the length of the study (Fig. 4 A, C & E). The only other mosquito species collected within the bay island site was *Aedes alternans*, a large mosquito with predatory larvae (Webb et al., 2016). Container-associated mosquitoes, such as *Culex quinquefasciatus* (Say) and *Aedes notoscriptus* (Skuse), made up the majority of the remainder of residential collections. A minimal number of non-target insects were collected with similar body sizes and types to mosquitoes (e.g., non-biting *Chironomidae* midges) within each habitat, likely due to the lack of a light source on the trap or nearby artificial light sources. Even fewer (3.14 ± 1.14%) insects classified as ‘large’ by the BG-Counter were collected. The majority of ‘large’ classed insects belonged to the orders Lepidoptera, Diptera, Coleoptera, and Hymenoptera. Insects that would be classified as ‘small’ made up the second largest (35.48 ± 11.14%) group by counts of the three classifications (the other being mosquito-sized insects). This size class was dominated by *Culicoides* biting midges (mostly *C. ornatus*, *C. molestus*, and *C. marmoratus*).

**Figure 3.**
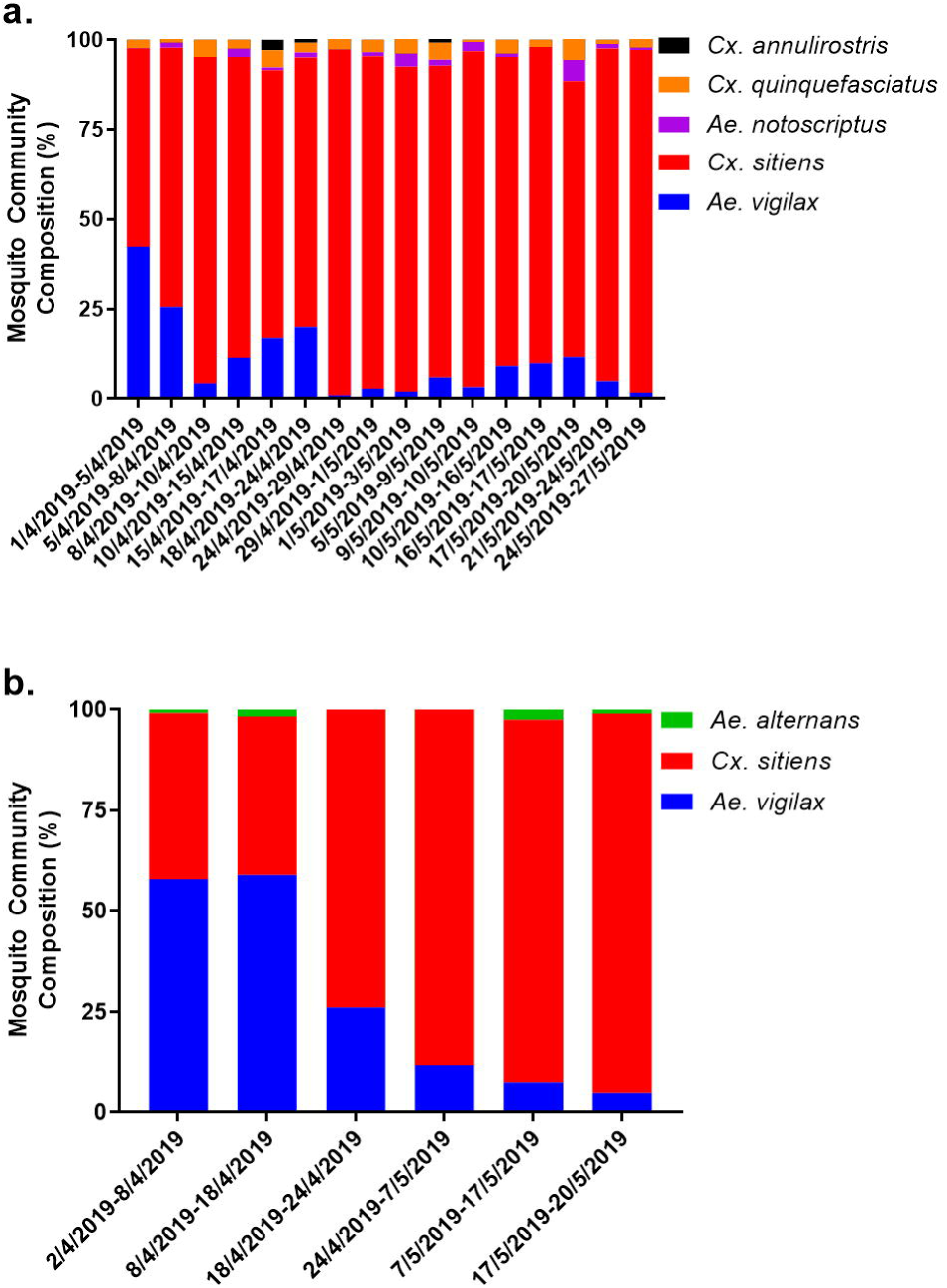
The relative occurrence of mosquito species within a (A) residential coastal development and (B) adjacent uninhabited bay island during the study period. Saltmarsh species include *Ae. vigilax*, *Cx. sitiens* and *Ae. alternans*.

**Figure 4.**
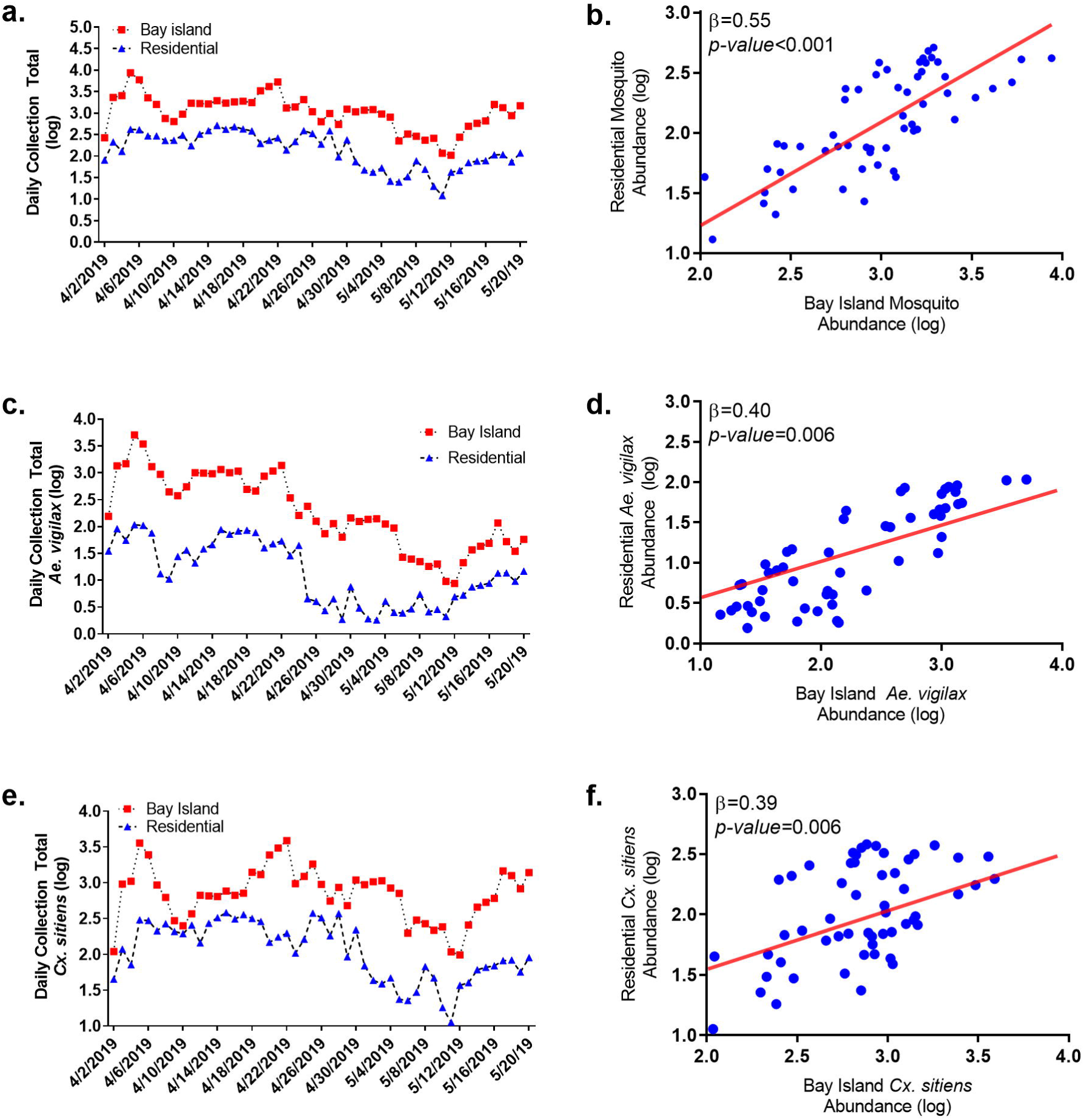
Daily comparisons of saltmarsh mosquito abundance within a residential coastal development and adjacent uninhabited bay island, including (A-B) total saltmarsh mosquito abundance, (C-D) estimated *Ae. vigilax* abundance, and (E-F) estimated *Cx. sitiens* abundance. β is the regression coefficient value for each predictor (bay island source abundance) of residential abundance determined by generalized least squares regression. The red line represents the best-fit line of the regression.

### 3.2 Agreement Rates between Manual and Automated Mosquito Counts

The mean agreement rate (Table 1) between manual and automated mosquito counts was 85.12% (95% CI: 77.36 - 92.89 %). The inclusion of ‘small’ classed insects tended to overestimate the number of mosquitoes (117.3 %; 95% CI: 88.07-146.5), whereas the inclusion of ‘large’ classed insects tended to underestimate the number of mosquitoes (80.84 %; 95% CI: 70.97-90.72 %). These observations indicate the use of ‘mosquito’ classed insect counts as the most representative of actual abundances and demonstrate that the BG-Counter can reliably and accurately differentiate target mosquitoes from other insects.

### 3.3 Synchronicity in Adult Mosquito Abundance

Linear regression analysis revealed a strong, positive relationship between bay island and residential daily mosquito abundances (Fig. 4 B, D & F; Table S1). The strongest predictor of daily residential mosquito abundance was daily abundance on the bay island (β=0.55±0.11, *t*=4.87, *p-*value<0.001), a relationship that held when applied to estimated *Ae. vigilax* (β=0.40±0.14, *t*=2.89, *p-*value=0.006) and *Cx. sitiens* (β=0.39±0.14, *t*=2.88, *p-*value=0.006) abundances. The strength of this relationship only worsened if additional time lags were applied (1-7 d). All other predictors were found to be insignificant.

### 3.4 Daily Host-Seeking Activity of Residential and Bay Island Populations

Patterns of daily host-seeking activity were similar between the bay island and residential sites with no significant (*F*_1,879_=0.47, *p=*0.50) differences observed within the defined 3 hr activity time periods (Fig. 5 A; Table S2). In general, host-seeking activity was highest between the hours of 18:00 to 06:00 (6 pm-6 am), with the greatest peak between 18:00 to 21:00 (6 pm-9 pm), and the lowest level of activity from 06:00 to 15:00 (3 am-3 pm), a pattern observed within both locations. A surprisingly high and constant level of host-seeking activity was observed between 21:00 to 03:00 (9 pm-3 am), a period not typically associated with high activity in crepuscular species (Reisen et al., 1976). High periods of host-seeking activity were associated with decreases in light intensity, increases in relative humidity, and decreases in ambient temperature (Fig. 5 B-D). Although significant differences in these variables were observed between the two sites, these differences occurred during daylight hours when there was little to no host-seeking activity. Of the three variables, light intensity was the strongest indicator of activity, with activity occurring immediately after sunset and ceasing immediately before sunrise (Fig. 6). The residential site had extensive environmental shading (i.e., house overhangs and larger tree cover) and subsequently recorded substantially lower levels of light intensity during daylight (6 am-6 pm) hours (mean = 2.74 ± 0.93 vs. 17.47 ± 4.89 lumens). Environmental shading also likely attributed to the higher levels of humidity (mean = 81.85 ± 2.71% vs. 66.06 ± 3.72%) and lower temperatures (mean = 22.82 ± 1.04 vs. 24.13 ± 1.36) recorded in the residential site relative to those recorded on the bay island.

**Figure 5.**
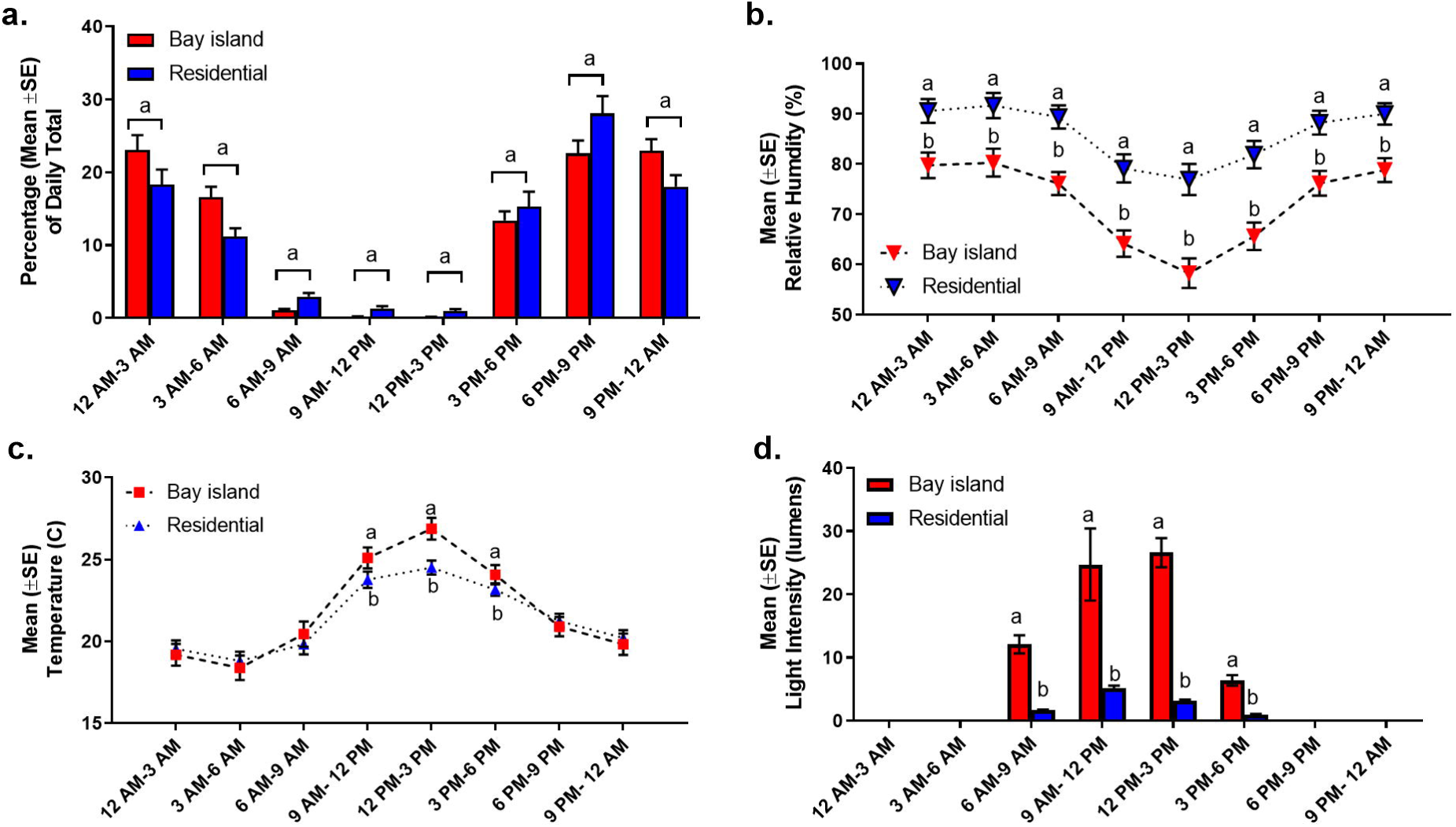
Comparisons of saltmarsh (A) mosquito host-seeking and recorded environmental variables (B. relative humidity; C. temperature; and D. light intensity) within a residential coastal development and adjacent uninhabited bay island. Points that do not share the share similar letters denote statistical significance (*p-value*<0.05 two-way ANOVA). All values are means ± SE of daily observations during the specified time periods.

**Figure 6.**
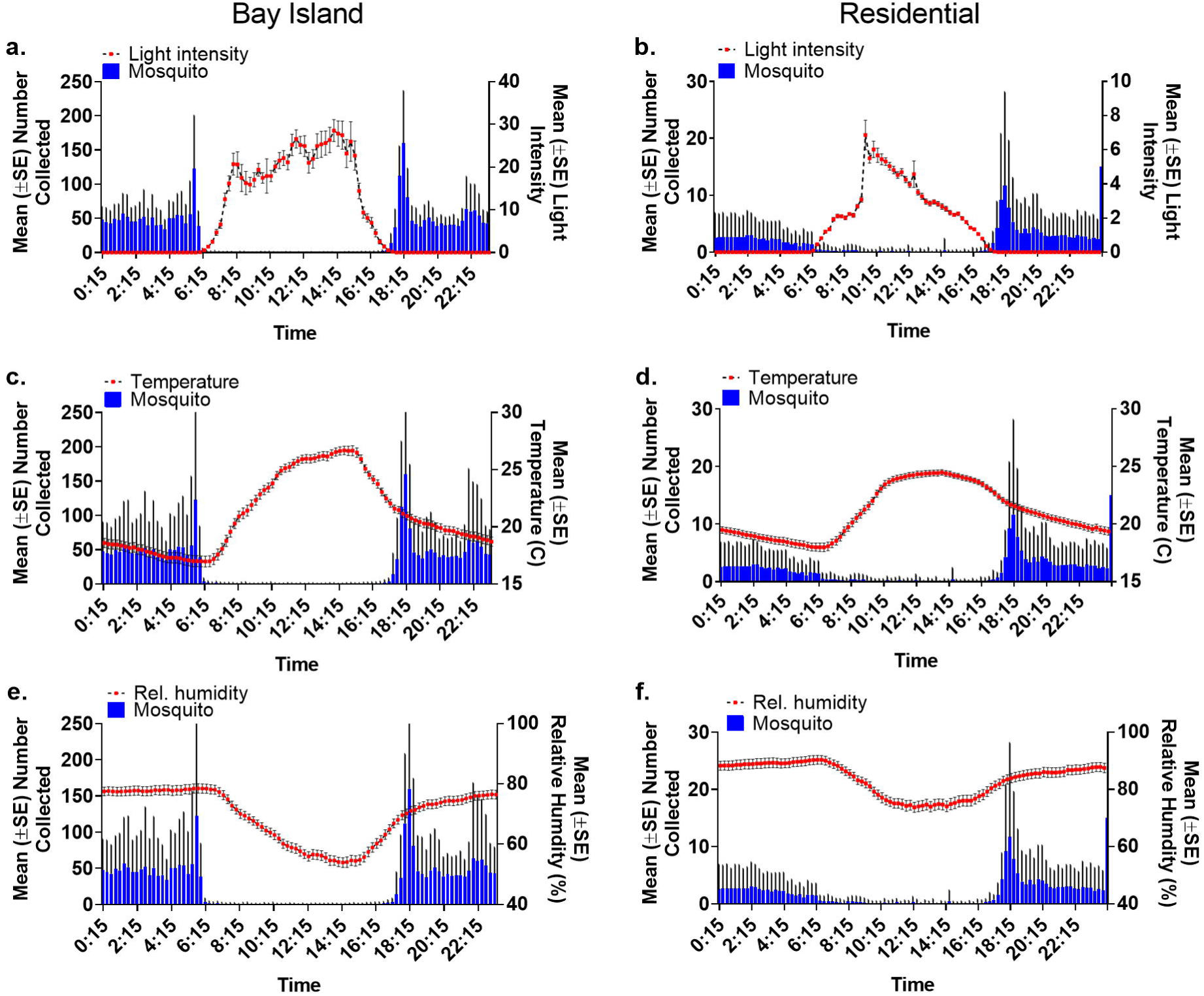
Daily trends in saltmarsh mosquito host-seeking activity and recorded environmental variables (A-B light intensity; B-C temperature; and E-F relative humidity) within a coastal development and adjacent uninhabited bay island. All values are means ± SE of individual 15-minute observation intervals.

## 4. Discussion

Mosquito nuisance and the threat of disease is given little consideration in urban planning despite considerable research on the topic and substantial costs to both individuals and government (Dale, 2010; Dwyer et al., 2016; Irwin et al., 2008; Medlock and Vaux, 2015). Instead, regional planning guidelines promote the incorporation of thinly supported mitigation strategies such as the use of green space *buffer zones* that are likely of little use against highly vagile mosquitoes (Dwyer et al., 2016; Webb and Russell, 2019). The results of the present study provide further evidence against the efficacy of such buffers by highlighting the dispersal of nuisance mosquitoes from a remote offshore breeding site to a coastal development over a considerable expanse (ca. 1.4 km) of open water. The nature of this dispersal was unidirectional and driven solely by island population dynamics. We must, however, acknowledge that dispersal of mosquitoes from neighboring breeding sites was likely and may have influenced observed trends. Nevertheless, the proximity of the two study sites and large area of productive mosquito habitat within the bay island, combined with a lack of notable onshore breeding sites and unfavorable nightly winds, suggest that the influence of such dispersal was minimal. Thus, our results demonstrate that dispersal of mosquitoes from distant saltmarsh habitats, including offshore breeding sites, are likely to be primary sources of mosquito nuisance and disease risk for coastal housing developments. This observation underscores the need to develop new planning and regulatory guidelines that alert urban planners to the risks of encroaching on habitats close to the sources of highly vagile mosquito species as well as to the impracticalities of currently accepted mitigation strategies.

Effectively controlling highly vagile mosquito species is no small task. In southeast Queensland, the control of *Ae. vigilax* is the focus of area-wide larviciding campaigns largely reliant on the use of the biological agent *Bacillus thuringiensis israelensis* (*Bti*). Outside of habitat modifications, such as runnelling and open water management (Dale and Breitfuss, 2009; Dale and Knight, 2006), the targeted application of *Bti* is widely accepted as the most effective means of control (Becker, 2006; Dale and Hulsman, 1990; Floore, 2006). However, inundating tides, dense mangrove canopy, and the occurrence of staggered broods (non-synchronous egg hatching) makes complete control challenging. Many local authorities have therefore advocated for the incorporation of an area of cleared green space between housing developments and known mosquito habitats, commonly referred to as a buffer zone (Webb and Russell, 2019), to minimize contact between mosquitoes and local residents. The use of buffer zones is founded on the principal that a provision of land surrounding residential developments devoid of mosquito refuge will minimize the dispersal of mosquitoes from nearby breeding sites into adjacent developments. Despite the inclusion of buffer zones in many local and state mosquito management plans (Dwyer et al., 2016; Scott, 2002), the practice is based on a rudimentary understanding of the factors influencing the spatial and temporal dispersal of a handful of pest mosquitoes. Although buffer zones may reduce the hazard from species that disperse short distances (<1 km), they are probably of little use against highly vagile species such as those studied here. A redrafting of local and regional regulatory guidelines is therefore needed. Efforts should be made to ensure that all future guidelines incorporate a thorough understanding of the dispersal capabilities, biting behaviour and relative abundances of key mosquito species.

The fine-scale patterns of population synchronicity observed in this study has additional operational significance as it appears to refute previous assumptions of *Ae. vigilax* dispersal. The high rate of autogeny (71-100%) in this species (Hugo et al., 2003) has long been thought to postpone dispersal resulting in delayed human contact (Reisen and Milby, 1987; Spadoni et al., 1974). The data collected in this study suggests this is an erroneous assumption, a conclusion further supported by other studies linking residential nuisance complaints to increases in offshore populations (Ryan et al., 2006). Although most vector-borne disease transmission cycles are complex and the density of vector mosquitoes is not always correlated with pathogen transmission intensity (Beier et al., 1999), earlier studies in the same region have shown that mainland areas most at risk of experiencing significant incursions of offshore *Ae. vigilax* have higher than expected numbers of RRV disease cases (Ryan et al., 2006). This observation gives additional credence to the need for improved area-wide control strategies that can overcome the operational limitations imposed by saltmarsh environments. Importantly, the synchronicity in host-seeking activity observed indicates that site-specific environmental differences have little impact on the biting activity of *Ae. vigilax* and *Cx. sitiens* dominated communities. This is in contrast to seasonally-driven differences that are known to significantly alter host-seeking activity (Chadee and Corbet, 1987; Jaenson, 1988; Klein et al., 1992). This suggests that adult control activities targeting these species can be synchronized across habitats to maximize control, but it also highlights the need for long-term studies so that both residents and control programs can alter their behaviours or control strategies in response to seasonal variations in host-seeking activity.

While the disease risks associated with mosquito abundance have been extensively studied in Australia and elsewhere (Hu et al., 2006; Ryan et al., 2006; Ryan et al., 1999; Whelan et al., 2003), the impact of mosquito nuisance on residents’ quality of life is rarely investigated (Curco et al., 2008; Halasa et al., 2014; Ratigan, 2000). Such information is essential to guiding local and regional planning decisions. The personal costs of mosquito nuisance are considerable with many homeowners placing more importance on nuisance than disease in terms of their demand and willingness to pay for mosquito control (Dickinson and Paskewitz, 2012; von Hirsch and Becker, 2009). In some cases, the impact of mosquito nuisance on resident quality of life has been equated to the loss from a worrisome health problem resulting in upwards of 80% of residents opting to stay indoors to avoid mosquitoes (Halasa et al., 2014; Ratigan, 2000). This underscores the importance of understanding the health, economic and personal costs of mosquito nuisance when developing local and regional planning guidelines. Steps should be taken to foster a greater public understanding of the mosquito species of concern and the risks they pose, and what residents can do to mitigate these risks. This may include the sharing of routine trapping data, population forecasting, and daily and seasonal patterns of biting activity by local government agencies and developers.

Lastly, it is important to note the lack of a detectable relationship between local wind patterns and changes in the studied mosquito populations. Insects often achieve long-distance dispersal by utilising the rapid winds situated above their flight boundary layer, a layer of atmosphere usually close to the ground where wind speeds are low enough to facilitate insect flight (Srygley and Dudley, 2008; Taylor, 1974). A lack of impact of wind direction on abundance in this study is therefore remarkable. Although the region experienced winds favourable to mainland dispersal, they occurred predominantly during daytime hours when mosquitoes were least active and likely sheltering in low-lying vegetation. As such, few mosquitoes would be expected to be captured by daytime winds. In contrast, winds were largely unfavorable to mainland dispersal during the period of greatest mosquito activity (i.e., late evening to early morning). Observed patterns of population synchronicity between the two studied habitats demonstrates considerable directed dispersal by both *Ae. vigilax* and *Cx. sitiens* in spite of unfavourable wind patterns.

## 5. Conclusions

When planning new coastal developments, it is imperative to consider the pest, public health and quality of life impacts of mosquitoes dispersing from often overlooked distant breeding sites. This is especially the case when highly vagile saltmarsh species are the primary concern. Local authorities are increasingly supporting changes to urban planning guidelines to mitigate the risks posed by mosquitoes, yet existing strategies such as buffer zones around residential developments are likely to have little impact against highly vagile species like those studied here. Such strategies are also impractical in existing urbanized coastal waterfronts where they simply cannot not be implemented retrospectively. Consequently, new regulatory guidelines are needed that incorporate a thorough understanding of the dispersal capabilities and relative abundances of key mosquito species. Such guidelines should include a proviso that residents be warned of the risks of mosquito nuisance to their health and lifestyle. Emerging smart trap technologies can assist these efforts by generating data on mosquito abundance, host-seeking activity and their relationships to local environmental factors at levels previously unattainable.

## Acknowledgements

We thank the staff of the Redland City Council Pest Management team for their assistance with trapping mosquitoes and for providing transport to field sites. The Mosquito and Arbovirus Research Committee (Inc.), an independent Australian body involved in mosquito research made up of representatives from local governments, state government agencies, industry and scientific organisations, provided the funding for the study.

