## Supplementary material for "What happens on islands, doesn’t stay on islands: Patterns of synchronicity in mosquito nuisance and host-seeking activity between a mangrove island and adjacent coastal development": Table S1

**Appendices**

**Table A1.** Summary of generalized least squares regression analysis.

| Predictors of residential abundance (total) | | | | |  |
| --- | --- | --- | --- | --- | --- |
| Coefficient | Value | Std. Error | t-value | p-value | Sign. |
| Bay island (total) mosquito abundance | 0.55 | 0.11 | 4.87 | <0.001 | Yes |
| Wind speed night | -0.018 | 0.024 | -0.74 | 0.46 | No |
| Wind speed day | 0.018 | 0.015 | 1.24 | 0.22 | No |
| Wind direction | -0.0005 | 0.0009 | -0.55 | 0.58 | No |
| Daily mean temperature | 0.036 | 0.018 | 1.89 | 0.07 | No |
| Bay island mosquito abundance : wind speed night | -0.029 | 0.08 | -0.37 | 0.71 | No |
| Bay island mosquito abundance : wind speed day | 0.022 | 0.067 | 0.32 | 0.75 | No |
| Bay island mosquito abundance : wind direction | -0.002 | 0.002 | -0.78 | 0.44 | No |
| Predictors of residential *Aedes vigilax* abundance | | | | |  |
| Coefficient | Value | Std. Error | t-value | p-value | Sign. |
| Bay island *Ae. vigilax* abundance | 0.4 | 0.14 | 2.89 | 0.006 | Yes |
| Wind speed night | 0.04 | 0.028 | 1.33 | 0.19 | No |
| Wind speed day | 0.004 | 0.018 | 0.24 | 0.81 | No |
| Wind direction | -0.0005 | 0.0011 | -0.48 | 0.63 | No |
| Daily mean temperature | 0.02 | 0.02 | 0.93 | 0.36 | No |
| Bay island *Ae. vigilax* abundance : wind speed night | 0.03 | 0.047 | 0.64 | 0.53 | No |
| Bay island *Ae. vigilax* abundance : wind speed day | -0.04 | 0.04 | -0.99 | 0.32 | No |
| Bay island *Ae. vigilax* abundance : wind direction | -0.016 | 0.042 | -0.39 | 0.7 | No |
| Predictors of residential *Culex sitiens* abundance | | | | |  |
| Coefficient | Value | Std. Error | t-value | p-value | Sign. |
| Bay island *Cx. sitiens* abundance | 0.39 | 0.14 | 2.88 | 0.006 | Yes |
| wind speed night | -0.006 | 0.02 | -0.29 | 0.77 | No |
| wind speed day | 0.062 | 0.025 | 2.51 | 0.016 | No |
| wind direction | 0.005 | 0.011 | 0.47 | 0.64 | No |
| Daily mean temperature | 0.019 | 0.019 | 0.99 | 0.32 | No |
| Bay island *Cx. sitiens* abundance : wind speed night | 0.005 | 0.073 | 0.079 | 0.94 | No |
| Bay island *Cx. sitiens* abundance : wind speed day | -0.025 | 0.068 | -0.36 | 0.72 | No |
| Bay island *Cx. sitiens* abundance : wind direction | -0.001 | 0.004 | -0.34 | 0.73 | No |

**Table A2.** Summary of recorded environmental variables during each host-seeking activity period.

|  | Temperature (^o^C) | | | | | |
| --- | --- | --- | --- | --- | --- | --- |
|  | Bay island | | Residential | |  |  |
| Time | Mean | SE | Mean | SE | Difference Significant | Adjusted P Value |
| 12 AM-3 AM | 19.19 | 0.33 | 19.56 | 0.25 | No | 0.97 |
| 3 AM-6 AM | 18.39 | 0.36 | 18.82 | 0.27 | No | 0.95 |
| 6 AM-9 AM | 20.45 | 0.38 | 19.83 | 0.30 | No | 0.67 |
| 9 AM- 12 PM | 25.10 | 0.31 | 23.78 | 0.24 | Yes | 0.01 |
| 12 PM-3 PM | 26.87 | 0.33 | 24.51 | 0.21 | Yes | <0.001 |
| 3 PM-6 PM | 24.08 | 0.29 | 23.16 | 0.19 | No | 0.18 |
| 6 PM-9 PM | 20.90 | 0.29 | 21.24 | 0.22 | No | 0.98 |
| 9 PM- 12 AM | 19.83 | 0.32 | 20.21 | 0.24 | No | 0.96 |
|  | Light Intensity (lumens) | | | | | |
|  | Bay island | | Residential | |  |  |
| Time | Mean | SE | Mean | SE | Difference Significant | Adjusted P Value |
| 12 AM-3 AM | 0 | 0 | 0 | 0 | No | 0.99 |
| 3 AM-6 AM | 0.0136 | 0.0044 | 0 | 0 | No | 0.99 |
| 6 AM-9 AM | 12.121 | 0.7157 | 1.6447 | 0.0619 | Yes | <0.001 |
| 9 AM- 12 PM | 24.724 | 2.8397 | 5.1929 | 0.1925 | Yes | <0.001 |
| 12 PM-3 PM | 26.618 | 1.1403 | 3.1574 | 0.1105 | Yes | <0.001 |
| 3 PM-6 PM | 6.4182 | 0.4197 | 0.9782 | 0.0463 | Yes | <0.001 |
| 6 PM-9 PM | 0 | 0 | 0 | 0 | No | 0.99 |
| 9 PM- 12 AM | 0 | 0 | 0 | 0 | No | 0.99 |
|  | Relative Humidity (%) | | | | | |
|  | Bay island | | Residential | |  |  |
| Time | Mean | SE | Mean | SE | Difference Significant | Adjusted P Value |
| 12 AM-3 AM | 79.766 | 1.2587 | 90.607 | 1.187 | Yes | <0.001 |
| 3 AM-6 AM | 80.274 | 1.3783 | 91.698 | 1.2463 | Yes | <0.001 |
| 6 AM-9 AM | 76.136 | 1.1469 | 89.404 | 1.1491 | Yes | <0.001 |
| 9 AM- 12 PM | 64.177 | 1.311 | 79.148 | 1.389 | Yes | <0.001 |
| 12 PM-3 PM | 58.286 | 1.4612 | 76.952 | 1.5394 | Yes | <0.001 |
| 3 PM-6 PM | 65.656 | 1.3679 | 81.9 | 1.3549 | Yes | <0.001 |
| 6 PM-9 PM | 76.187 | 1.2219 | 88.273 | 1.1832 | Yes | <0.001 |
| 9 PM- 12 AM | 78.821 | 1.1789 | 90.014 | 1.0536 | Yes | <0.001 |
